# Sex bias determines MERS-CoV infection outcomes in a mouse model of differential pathogenicity

**DOI:** 10.1101/2025.06.19.660369

**Authors:** Marin R. Habbick, Trenton Bushmaker, Reema Singh, Pryanka Sharma, Vincent J. Munster, Neeltje van Doremalen, Angela L. Rasmussen

**Affiliations:** Vaccine and Infectious Disease Organization, University of Saskatchewan, Saskatoon, SK, Canada; Department of Biochemistry, Microbiology, and Immunology, University of Saskatchewan, Saskatoon, SK, Canada; Laboratory of Virology, Division of Intramural Research, National Institute of Allergy and Infectious Diseases, National Institutes of Health, Rocky Mountain Laboratories, Hamilton, MT, USA; Center for Infection and Immunity, Columbia Mailman School of Public Health, New York, NY, USA; Department of Ecology and Evolution, Stony Brook University, Stony Brook, NY, USA

## Abstract

Middle East respiratory syndrome coronavirus (MERS-CoV) causes a spectrum of disease outcomes in infected humans, ranging from asymptomatic or tolerant to lethal. While the virus itself contributes to pathogenesis, disease severity is primarily influenced by the host’s response to infection. One factor observed to impact the host response is sex, as epidemiological data indicates that male persons have a higher case fatality rate than females infected with MERS-CoV. However, the mechanism underlying this sex bias is unknown and disease course remains difficult to predict. This study investigates how male and female transgenic mice expressing humanized dipeptidyl peptidase-4 (hDPP4) respond to MERS-CoV infection following exposure to either a tolerance-inducing low dose or lethal high dose. We observed that female hDPP4 mice display dose-dependent tolerance to infection and males experienced uniformly lethal disease in both dosing groups. Longitudinal transcriptomic analysis revealed that males suppress innate and inflammatory responses early after infection, causing delayed induction of the host antiviral response. In contrast, high dose females mount an immediate and sustained interferon and inflammatory response, activating antiviral effectors and interferon-stimulated genes. Tolerant females displayed the greatest transcriptional control, showing no pathway enrichment and minimal changes in weight throughout infection. Our results suggest that the magnitude of the response is driven by dose while the nature of the response in shaped by sex. Females mount a more robust response to MERS-CoV infection, allowing females to tolerate low-dose infection but causing uncontrolled inflammation after high dose infection. In contrast, males experienced lethal outcomes regardless of dose. By examining the dynamics of sex-biased host transcriptional responses in determining disease severity, this study highlights the importance of sex as a biological variable in coronavirus pathogenesis research.

## Introduction

Middle East respiratory syndrome coronavirus (MERS-CoV) is the most lethal human CoV among diagnosed cases that has emerged to date (WHO 2022). With a case fatality rate of approximately 35% in severe cases, ongoing zoonotic transmission and no available treatments or vaccines, MERS-CoV has continued to pose a significant threat to public health since it emerged in Saudi Arabia in 2012 (Zaki et al. 2012; Kelly-Cirino et al. 2019) . MERS-CoV outbreaks usually begin through zoonotic transmission from dromedary camels and can result in subsequent nosocomial transmission (Azhar et al. 2014; Hui, Perlman, and Zumla 2015; Assiri et al. 2013). As of May 2024, more than 2,500 laboratory-confirmed cases of MERS-CoV have been identified across 27 countries leading to 943 deaths (WHO 2024b); however, the true case number is likely higher, as mild cases of MERS-CoV often go unreported (WHO 2022).

Different hosts infected with the same virus may exhibit a range of clinical disease phenotypes, from mild to lethal illness (Soares, Teixeira, and Moita 2017). Some individuals display tolerance to MERS-CoV infection, meaning they are productively infected and shed virus, but display subclinical or no clinical signs of disease (Soares, Teixeira, and Moita 2017). Tolerant hosts mount a tightly controlled immune response, upregulating then downregulating the inflammatory response to efficiently clear virus without damaging infected tissues (Flerlage et al. 2021; Soares, Teixeira, and Moita 2017). Alternatively, hosts that mount a dysregulated immune response may experience moderate or severe infection as the virus is inefficiently cleared, resulting in an overactive and prolonged inflammatory response that damages lung tissue (Lopes-Pacheco et al. 2021; Soares, Teixeira, and Moita 2017; Zhu et al. 2020). Although the difference in MERS-CoV pathogenicity between a tolerant and severe outcome is in part due to the action of the virus itself, a major determinant of disease severity is the host response (Widagdo et al. 2019; Memish et al. 2020; Prescott et al. 2018; Falzarano et al. 2014; de Wit et al. 2013). Certain risk factors, such as old age and specific underlying comorbidities, are known to increase the likelihood of severe disease (Memish et al. 2020); however, pathogenicity remains difficult to predict. Another poorly understood risk factor that may influence disease severity is sex, as epidemiological data indicates that male persons have a higher case fatality rate than females (Alghamdi et al. 2014). Differential sex hormone signalling may contribute to this observed sex bias; however, the mechanisms underlying sex hormone action in the context of MERS-CoV infection have yet to be investigated.

To investigate the differences underlying a tolerant versus severe response to MERS-CoV infection, male and female transgenic Balb/c mice expressing human dipeptidyl peptidase 4 (hDPP4) were infected with either a low or high dose of MERS-CoV. Rodents are naturally resistant to MERS-CoV infection due to structural differences in murine DPP4, but can be rendered susceptible through transgenic expression of the human ortholog (Coleman et al. 2014; Li and McCray 2020). We collected clinical and viral titre data and confirmed that hDPP4 mice infected with a low or high dose of MERS-CoV are tolerant to infection or succumb to lethal disease, respectively. To study the host response dynamics of these differential outcomes, we performed longitudinal transcriptomic sampling. We found that mice with lethal disease mount early potent inflammatory responses to infection compared to the more tightly regulated responses observed in the tolerant group. In addition to investigating the dose-dependent response to infection, we disaggregated male and female samples to assess whether sex influences disease outcomes. Our analysis revealed a sex-biased response to infection, underscored by differences in timing and magnitude of the antiviral response between high dose males and females, as well as true tolerance observed in low dose females but not in low dose males. Distinct sex-specific gene expression signatures were linked to upstream sex hormone signaling. This aligns with observed sex biases in human cases of MERS-CoV and indicates potential mechanisms determining sex-specific disease severity and outcome.

## Results

### hDPP4 mice exhibit dose-dependent outcomes to MERS-CoV infection

To investigate differences in morbidity and survival, hDPP4 mice were infected with either a high (10^5^ median tissue culture infectious dose; TCID_50_) or low (10^2^ TCID_50_) dose of MERS-CoV, respectively (Fig. 1). Morbidity and mortality were observed daily for 15 days post-infection (dpi). For the first two days of infection, the weight of the mice in both dose groups remained relatively stable (Fig. 1A). By 3 dpi, significant differences were observed, lethal mice rapidly began to lose weight and succumbed to infection by 6 dpi (Fig. 1B). By day 7, 50% of the tolerant mice reached end-point criteria for euthanasia (Fig. 1B), accompanied by mild fluctuations in mean weight throughout the infection (Fig 1A). Survivors in the tolerant group regained their weight by day 15 (Fig. 1A). Differences in survivorship were significant between lethal and tolerant groups (Fig 1B). Thus, mice infected with a high dose of MERS-CoV developed greater disease severity and experienced uniform lethality compared to those infected with a low dose.

**Figure 1:**
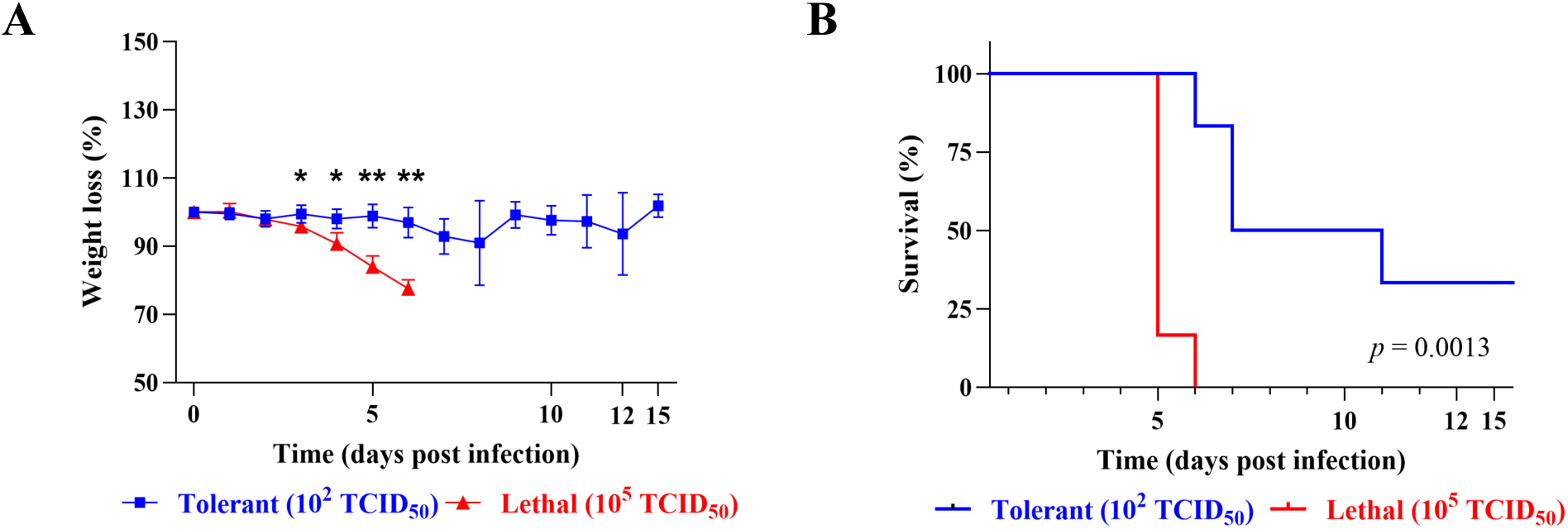
Distinct morbidity and mortality in transgenic hDPP4 mice infected with a high or low dose of MERS-CoV. Mice were infected at 10^5^ TCID_50_ or 10^2^ TCID_50_ or were mock-infected. **A)** Percent of starting body weight over the course of infection in high dose lethal (red) or low dose tolerant (blue) mice. Data shown are mean ± SEM from six mice per condition. Weights were compared between groups for each day using a multiple Mann-Whitney test with an adjusted 5% FDR to account for multiple comparisons (*\** = *p* < 0.05, ** = *p* < 0.01). **B)** Kaplan-Meier survival curve for lethal (red) and tolerant (blue) mice. Statistical differences between survival curves was determined using the log-rank test (*p* = 0.0013).

### Increased viral titre and load in lungs and brain of high dose mice

To validate the dose-dependent model of pathogenesis and assess differences in virus replication kinetics, mice were sacrificed on days 1-5 post-infection, and samples were collected to determine viral titre and load (Fig. 2). Lung samples were harvested for analysis by TCID_50_ as well as genomic and subgenomic RT-qPCR. Viral titre and load analysis showed higher levels of MERS-CoV infection in the lungs of lethally-infected mice on days 1-4 post-infection compared to the tolerant group, with statistically significant differences at 3 and 4 dpi (Fig 2A, 2C, S1A).

**Figure 2.**
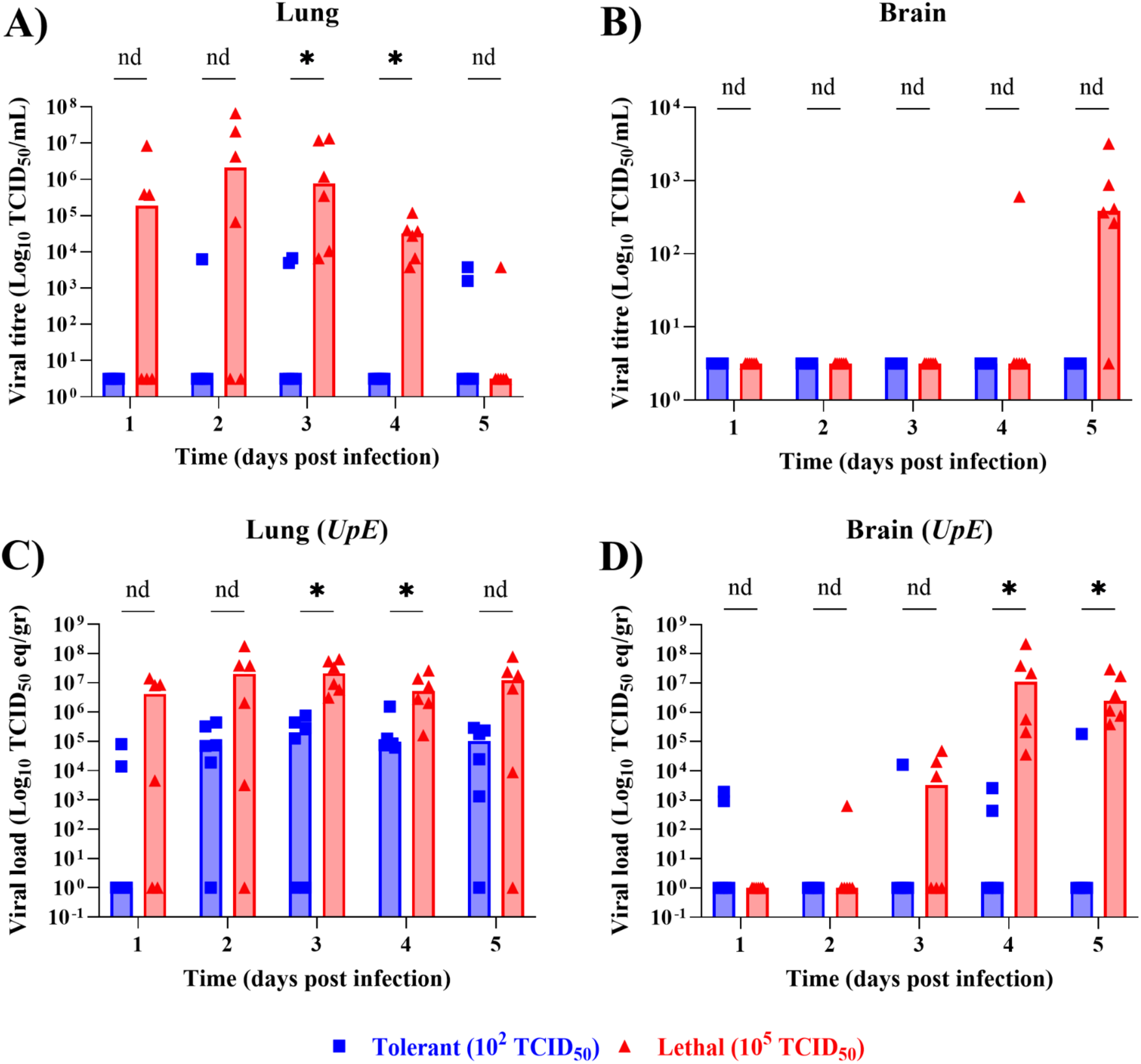
Mice infected with a 10^5^ TCID_50_ lethal dose of MERS-CoV exhibit increased viral replication compared to 10^2^ TCID_50_ group. Lung (**A, C**) and brain (**B, D**) tissues were collected days 1-5 post-infection and were subjected to virus titration (**A, B**) or RNA extraction (**C, D**). Infectious virus titre was assessed by TCID_50_ (**A, B**). Viral *UpE* genomic RNA was determined by RT-qPCR absolute quantitation in TCID_50_ equivalent/gram of tissue (TCID_50_ eq/gr) (**C, D**). Multiple Mann-Whitney t-tests were used to compare viral titre and load between lethal and tolerant groups with a 5% FDR adjusted *p-*value to account for multiple comparisons. Data are the median for 6 mice per group per timepoint. nd = not discoverable, * = *p* < 0.05.

In addition, brain samples were collected to investigate the abnormal encephalitis symptoms observed in transgenic mice expressing hDPP4 (Li et al. 2016). Unlike the tolerant group, lethal mice showed MERS-CoV replication in the brain and experienced increases in viral titre and load as infection progressed (Fig. 2B, 2D, S1B); however, significant differences were only observed for viral load at 4 and 5 dpi (Fig. 2D). Variations in viral titre and load in the brain between lethal and tolerant groups are dependent on initial infectious dose, suggesting that differential host responses are sufficient to control virus replication and extrapulmonary spread in the tolerant group.

### Lethal outcomes are associated with prolonged pathway enrichment

A major determinant of coronavirus disease severity is the host response. To investigate the response underlying tolerant versus severe outcomes to MERS-CoV infection, longitudinal RNA sequencing (RNA-seq) was performed on lung samples collected from 6 mice per dose group per timepoint. Samples were sequenced, normalized, counted and subjected to differential expression analysis (Fig. 3). Genes were determined to be differentially expressed (DE) if the false discovery rate-adjusted *p* < 0.05 and the fold change > |1.5| relative to time-matched mock-infected controls. To compare global gene expression profiles, we compared the number of genes meeting DE criteria (Fig. 3A, S2) and performed multidimensional scaling (MDS) for normalized read counts (S3) and DE genes (Fig. 3B). MDS is a dimensionality reduction method that allows high dimensional data, such as transcriptomic data, to be compressed into a 2 dimensional data point for comparing the similarity across samples. Differences in transcriptomes are indicated on an MDS plot by Euclidean distance between dimensionally compressed data points. The initial response to MERS-CoV infection at 1 dpi is similar between both dose groups (Fig. 3B), with the majority of DE genes downregulated (Fig. 3A). By 2 dpi, the response sharply diverges (Fig. 3B): the tolerant group has no genes meeting DE criteria, whereas the lethal group shifts to sustained upregulation of DE genes for the duration of infection (Fig 3A). The lethal group exhibits variability in their response to infection, with some samples clustering together and others overlapping with the tolerant group (Fig. 3B), indicating that some host responses are common to MERS-CoV infection independent of disease severity or outcome. In contrast, the tolerant group consistently clusters together across timepoints (Fig. 3B), transitioning from downregulation early in infection to upregulation by day 5 (Fig. 3A). The combination of heightened upregulation and increased heterogeneity in the lethal group may indicate a more dysregulated immune response compared to the more controlled trends observed in the tolerant group.

**Figure 3.**
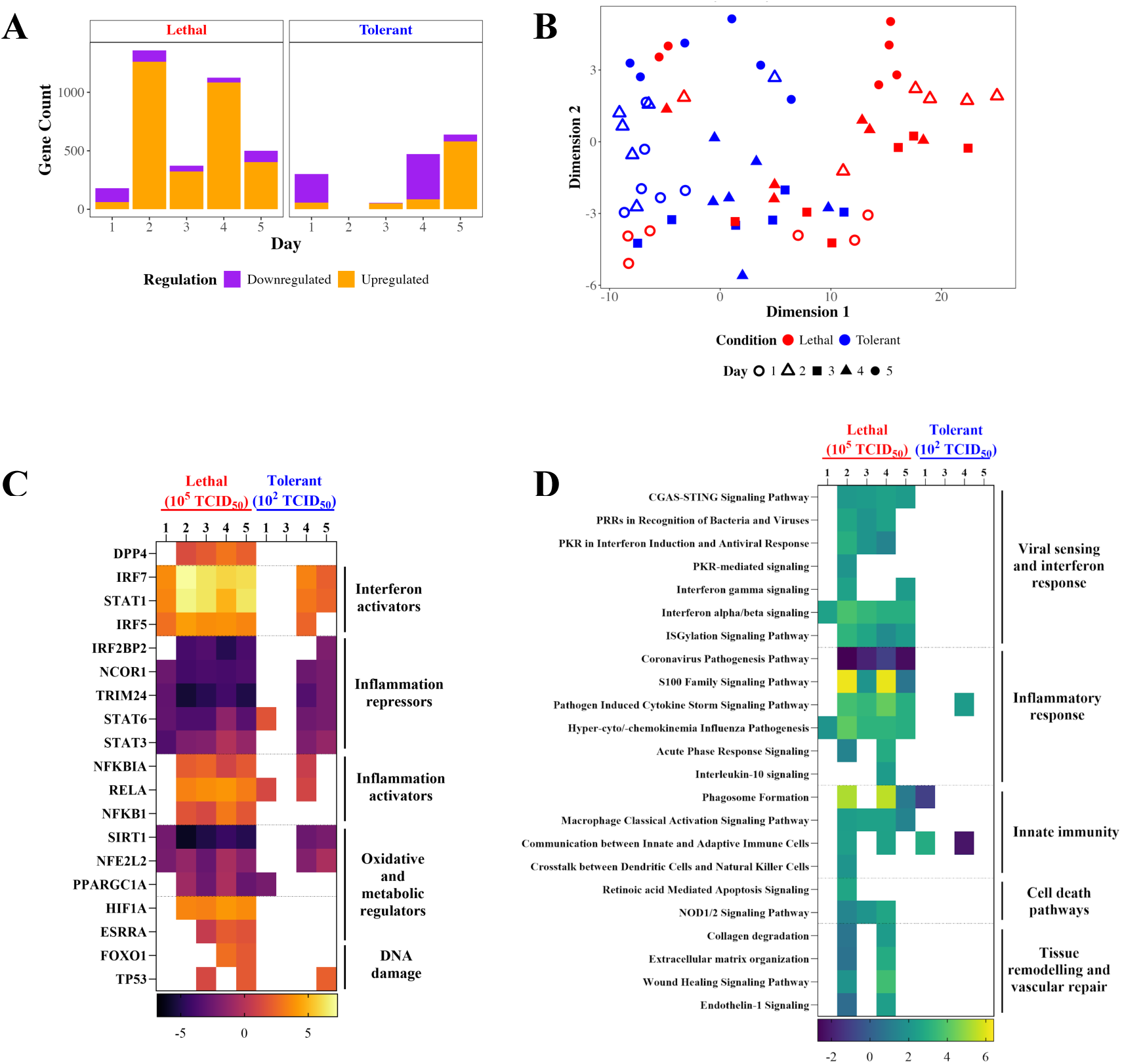
Low dose tolerant versus high dose lethal hDPP4 mice display differential pathway enrichment in response to MERS-CoV infection. hDPP4 mice were infected with MERS-CoV and lung samples were taken at days 1-5 post infection (N = 6 per time point per dose group). Samples were RNA-sequenced, filtered, annotated and analyzed for differential expression (DE; *p* < 0.05, log2FC > |1.5|). No genes met DE criteria for day 2 in the tolerant group. DE genes were visualized by bar-coded volcano plotting (**A**) and multidimensional scaling (**B**). DE genes were uploaded to Ingenuity Pathway Analysis (IPA) for functional analysis of upstream transcription factors (**C**) and canonical pathways (**D**) related to pathogenesis. IPA analyzes were conducted using a significance threshold of *p* < 0.01 and a Z-score ≥ |2|.

To interpret the biological relevance of these gene expression profiles, we performed functional analysis using Ingenuity Pathway Analysis (IPA), functional analysis software that assesses pathway enrichment and predicts pathway and gene activity using a manually curated functional knowledgebase derived from the peer-reviewed literature on mouse, rat, and human experimental systems (Fig. 3C, D). To define the MERS-CoV host response, DE genes uploaded to IPA were further filtered for Z-scores > |2| and BH-adjusted *p*-values < 0.01, selecting for upstream transcription factors (Fig. 3C; Data S1) and enriched pathways (Fig. 3D; Data S1) associated with pathogenesis. The tolerant group displays minimal predicted upstream transcriptional regulation (Fig. 3C) and pathway enrichment (Fig. 3D) compared to the lethal group, possibly indicating that low dose mice are better able to control infection, ameliorate pathogenicity, and maintain homeostatic gene expression. Conversely, the lethal group responds to infection immediately, upregulating transcriptional activators of the antiviral interferon (IFN) pathway and downregulating repressors of inflammation on day 1 post-infection (Fig. 3C). Early transcriptional events trigger a surge in inflammatory and innate immune signalling pathways, such as IFN, proinflammatory cytokine signaling, macrophage activation, and phagocytosis, at 2 dpi (Fig. 3D). As expected, hyperinflammation subsequently drives changes in predicted upstream transcription factor regulation associated with apoptosis and oxidative and metabolic stress (Fig. 3C), activating pathways associated with disease pathology (Fig. 3D). The tolerant group does not show signs of hyperinflammation, with delayed and weaker activation of innate immunity and no induction of inflammatory mediators until 4 dpi (Fig. 3C). Therefore, the tolerant group mounts a more tightly regulated response to infection, allowing for virus clearance without the uncontrolled inflammation leading to pathology observed in the lethal group.

### Female sex hormones likely play a greater role in shaping MERS-CoV outcomes in mice

Epidemiological evidence indicates that humans infected with MERS-CoV display sex-specific outcomes that may reflect underlying biased responses to infection (Alghamdi et al. 2014). To investigate the role of sex in determining MERS-CoV outcomes in the hDPP4 mouse model, transcriptomic infection data was disaggregated by sex and re-analyzed (Fig. 4). Because sex hormones are known to drive sex biased responses for other pathogenic betacoronaviruses, including SARS-CoV-1 and -2 (Stelzig et al. 2020; Abramenko et al. 2021; Mateus et al. 2022; Samuel et al. 2020; Tramontana et al. 2021; Channappanavar et al. 2017), general infection trends in sex hormone protein and receptor expression were assessed by upstream analysis (Fig. 4A). The female sex hormones, estrogens and progestogens, met the Z-score cut-off of > |2|, while the androgens did not. Different forms of estrogen were predicted to exert distinct upstream effects on the host response. Estriol is predicted to be downregulated during acute infection, while the main biologically active form of estrogen, beta-estradiol (E2), undergoes dynamic regulation. In lethal females, as well as in tolerant males and females, E2-dependent gene expression becomes strongly activated by day 4 or 5 post-infection. Lethal males also exhibit greater predicted E2 activation than tolerant males, although this occurred earlier at 2 dpi. Notably, males show a pronounced downregulation of E2 at 1 dpi. Predicted progesterone activity patterns are similar between males and females, whereas androgens are only predicted to regulate the response in males.

**Figure 4:**
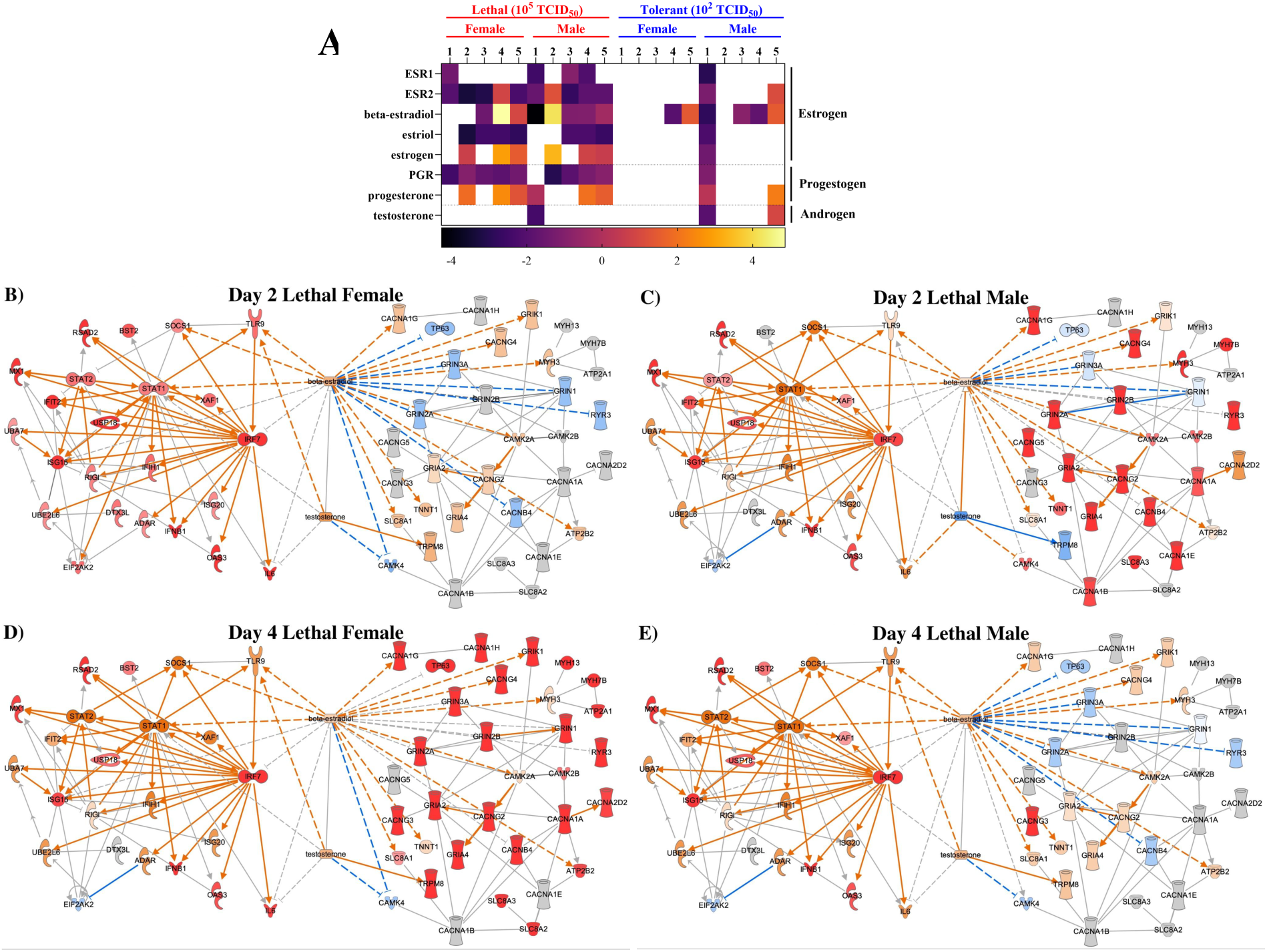
Estrogens may play a greater role than androgens in shaping MERS-CoV outcomes. Male (N = 2-4) and female (N = 2-4) mice were either mock-infected or infected with a lethal dose of 10⁵ median TCID_50_ or a lower, tolerance-simulating dose of 10² TCID_50_. On days 1-5 post-infection, RNA samples were collected from the lung and subjected to RNA-sequencing, data processing and differential expression (DE) analysis. DE genes (*p* < 0.05, log2FC > |1.5|) were uploaded to Ingenuity Pathway Analysis (IPA) for upstream sex hormone steroid and receptor analysis (A) and custom network mapping (B-E). The impact of estrogen and testosterone activity (middle) on antiviral interferon α/β and ISGylation DE molecules (left) were compared against pro-viral calcium signaling DE molecules (right). The IPA Overlay tool was used to visualize expression: red colouring denoted experimentally determined upregulation, while orange and blue represented predicted up or downregulation, respectively. Colour brightness indicated degree of enrichment. IPA upstream analyzes were performed using *p* < 0.01 and a Z-score ≥ |2| significance criteria. 2-4 male or female hDPP4 mice were used per time point per dose group.

To further investigate how sex hormones influence pathogenesis across interconnected host response pathways, we generated custom networks using IPA (Fig. 4B-I, Fig. S4A-D).

Specifically, networks were built by connecting DE molecules from heavily enriched antiviral pathways (interferon (IFN) α/β and ISGylation) and with calcium signaling, a heavily enriched proviral pathway implicated in facilitating viral entry, replication, and release (Chen, Cao, and Zhong 2019). Networks were generated at 2 and 4 dpi to capture the DE peaks observed in Figure 3A as well as host response dynamics relative to progression of infection and disease. Female and male mice exhibit temporal differences in their MERS-CoV response. At 2 dpi, lethal females strongly upregulate transcription of IFN α/β and ISGylation genes (Fig. 4B) while lethal males exhibit strong upregulation of calcium signaling molecules (Fig. 4C). By 4 dpi, lethal females and males show comparable antiviral enrichment; however, calcium signaling trends reverse: males no longer exhibit experimental enrichment (Fig. 4E) while calcium signaling becomes highly upregulated in females (Fig. 4D). Tolerant mice show minimal enrichment (Fig. S4); tolerant females are predicted to maintain anti-viral upregulation from 2 to 4 dpi (Fig. S4A, C), while tolerant males are only predicted to upregulate IFN α/β and ISGylation molecules by day 4 (Fig. S4B, D). Although the sex hormones show no experimentally-derived expression throughout infection, they display connections to both antiviral and calcium-related molecules. β-estradiol is connected directly or indirectly to more molecules in the network compared to testosterone, indicating that estrogens may exert a greater influence on determining host responses to MERS-CoV infection (Fig. 4B-4E). Overall, these findings suggest that female sex hormones play a more prominent role in shaping MERS-CoV infection outcomes compared to androgens.

### Female hDPP4 mice display dose-dependent MERS-CoV outcomes

To examine the role of sex in determining outcomes in the hDPP4 mouse model, morbidity and mortality data were disaggregated by sex and re-analyzed (Fig. 5). Females infected with a high dose of MERS-CoV exhibited uniform lethality by day 5 post-infection (Fig. 5B) with slightly greater but not statistically significant weight loss compared to males (Fig. 5A), who succumbed to infection by 6 dpi. The opposite is true of the tolerant group: males lost more weight compared to females, with uniform lethality observed by 11 dpi (Fig. 5B), while two of the three females survived and lost minimal weight (Fig. 5A). Statistically significant differences in weight were only observed between tolerant and lethal males or lethal females on days 5 and 6 post-infection (*p* < 0.05); however, with only 3 mice per sex per dose and mortality over time, the sample size is limited and must be interpreted with caution. Although survival did not differ between males and females within each dosing group, significant differences were observed between tolerant and lethal males (*p* < 0.05) as well as tolerant and lethal females (*p* < 0.05) (Fig. 5B).

**Figure 5.**
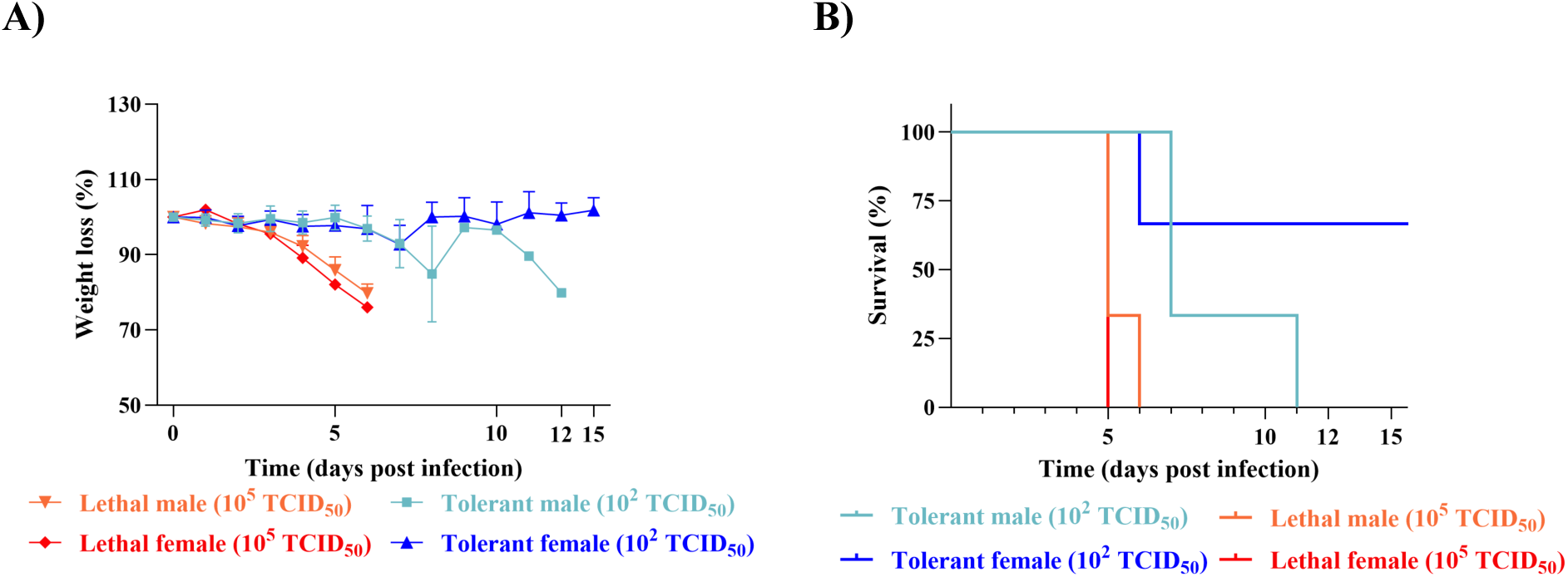
Male and female hDPP4 mice display sex differences in mortality and morbidity when infected with low or high dose MERS-CoV. Male and female mice were infected with a high (10^5^ TCID_50_) dose or low (10^2^ TCID_50_) dose to simulate tolerance or were mock-infected. **A**) Percent of starting body weight over the course of the infection in female high dose lethal (red, N= 3), male high dose lethal (orange, N= 3), female low dose tolerant (dark blue, N= 3) and male low dose tolerant (light blue, N= 3). Values represent mean ± SEM from three mice per condition. **B**) Kaplan-Meier survival curve for lethal female (red), lethal male (orange), tolerant female (dark blue) and tolerant male (light blue) mice. Statistical differences between survival curves were determined using the log-rank (Mantel-Cox) test. ns = not significant, * = *p* < 0.05.

### Female and male hDPP4 display sex-biased responses to MERS-CoV infection

To better understand how sex influences MERS-CoV outcomes, global differential gene expression trends and functional analyses were conducted by sex (Fig. 6). Genes meeting the DE cutoffs (log_2_ fold change > |1.5| and adjusted *p*-value < 0.05) relative to time-and sex-matched mock-infected controls were quantified (Fig. 6A, S5) and analyzed by MDS (Fig. 6B, S6) to comparatively examine transcriptomic trends. In addition to the dose-dependent differences observed previously (Fig. 3), sex-specific biases in DE gene expression were evident. Aside from 2 dpi, lethal females show greater transcriptional upregulation compared to lethal males (Fig. 6A). Both tolerant and lethal males exhibited greater transcriptional downregulation, especially at 1 dpi (Fig. 6A). However, low numbers of MERS-CoV read counts were observed in males on day 1 of infection (Fig. S7). Tolerant females demonstrated the lowest overall transcriptional response to infection across all groups (Fig. 6A). While some overlap between tolerant and lethal samples was observed within each sex, samples remained largely distinct, and with the exception of a single female sample from the lethal group on day 4, no overlap was observed between male and female responses (Fig. 6B).

**Figure 6.**
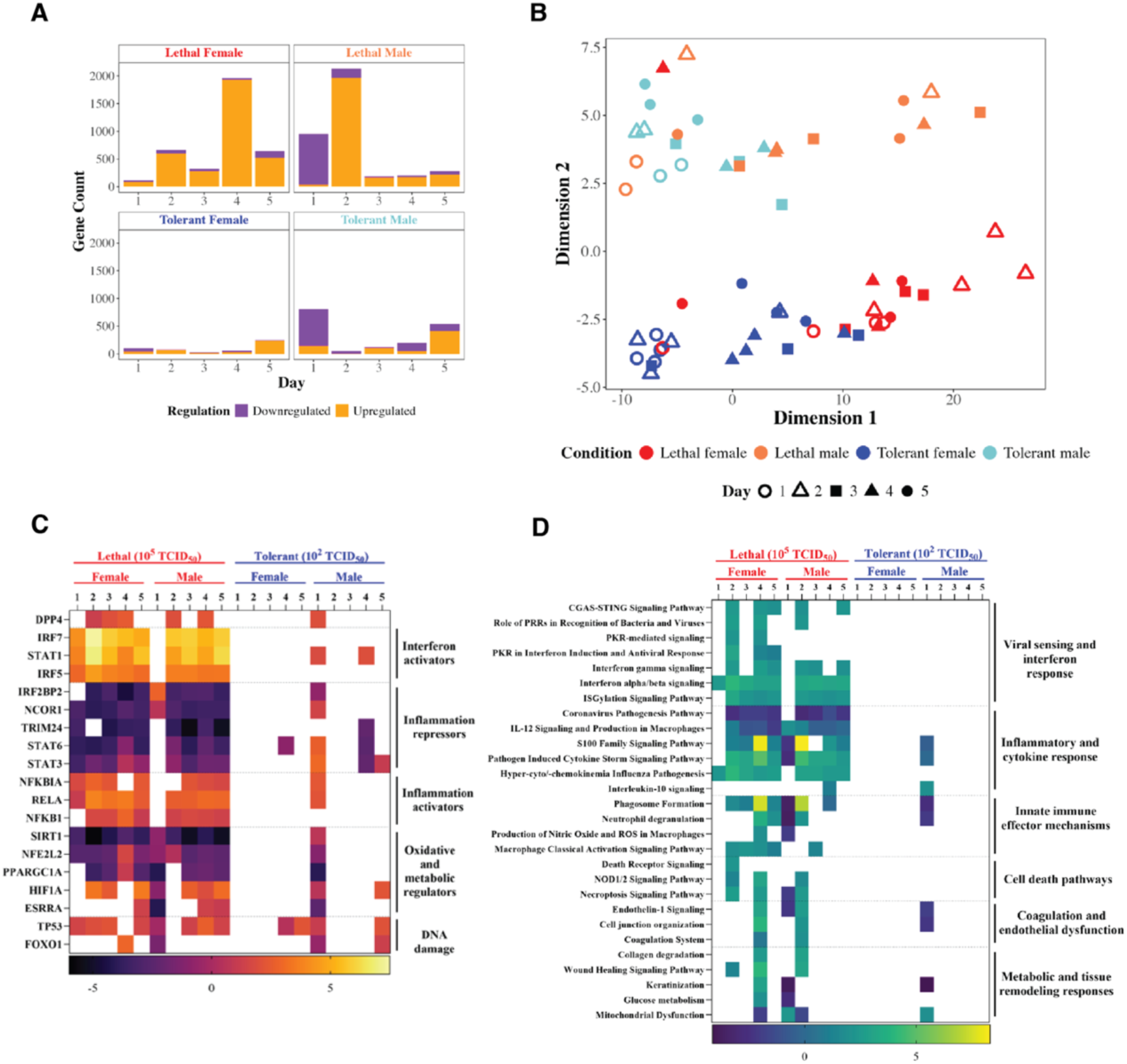
Male and female mice infected with MERS-CoV display both dose- and sex-dependent differences in their response to infection. Lung samples were collected from hDPP4 mice infected MERS-CoV on days 1-5 post-infection. Samples were subjected to RNA-seq followed by data processing and differential expression (DE) analysis. DE genes (*p* < 0.05, log2FC > |1.5|) were mapped using bar-coded volcano plotting (**A**) and multidimensional scaling (**B**). DE genes were uploaded to Ingenuity Pathway Analysis (IPA) to functionally analyze the impact of sex on upstream transcription factors (**C**) and canonical pathways (**D**) related to pathogenesis. IPA analyzes were conducted using a significance threshold of *p* < 0.01 and a Z-score ≥ |2|. 2-4 female or male hDPP4 mice were used per time point per dose group.

Finally, to assess how sex influenced pathways associated with viral pathogenesis, sex-specific functional analysis was performed (Fig. 6C, D). Upstream analysis and canonical pathways meeting previously described DE and functional enrichment criteria were investigated. Despite disaggregation by sex, the largest differences in enrichment were observed between tolerant and lethal groups; however, differences in upstream regulators (Fig. 6C; Data S2) and canonical pathways (Fig. 6D; Data S2) were still observed between males and females. Lethal female mice infected with MERS-CoV exhibit a robust and sustained antiviral and proinflammatory response, with enrichment of IFN α/β signalling and inflammatory pathways beginning on day 1 of infection. In contrast, lethal male mice show minimal antiviral activity on day 1, with no activation of viral sensing or IFN responses and inhibition of inflammation (Fig. 6D). Instead, lethal males show greater predicted activation of genes induced by upstream oxidative and metabolic regulators (Fig. 6C). However, the repression of the antiviral host response in males is not sustained, with lethal male mice showing strong upregulation of innate immune and inflammatory pathways by 2 dpi (Fig. 6D). In the lethal condition, both sexes activate pathogenesis-associated pathways–coagulation and endothelial dysfunction, metabolic and tissue remodelling, but males do so on day 2 compared to females on day 4 (Fig. 6D).

Pathway enrichment of protein kinase R (PKR)-related signalling and death-receptor signalling– antiviral effector mechanisms–was observed exclusively in high dose females (Fig. 6D). In contrast, sex-specific pathway enrichment in males was observed for IL-10 signalling, an anti-inflammatory cytokine influenced by androgen signalling (Fig. 6D) (Torcia et al. 2012). Tolerant male mice display a slight upregulation of IFN at 1 dpi, along with repression of inflammation, suggesting a more controlled immune response compared to lethal males (Fig. 6C, D). Unlike males in the low dose group, tolerant females survive MERS-CoV infection, displaying minimal upstream regulator prediction and no canonical pathway enrichment (Fig. 6C, D). Overall, these results suggest sex-specific differences in both the magnitude and timing of the antiviral response in hDPP4 mice during MERS-CoV infection.

## Discussion

This study provides novel insights into the mechanisms driving differential pathogenesis in MERS-CoV-infected hDPP4 mice. While previous research in this model has primarily focused on establishing susceptibility and modeling severe disease (Agrawal et al. 2015; Zhao et al. 2014; Li et al. 2016; Fan et al. 2018; Pascal et al. 2015; Cockrell et al. 2016; Li et al. 2017; Douglas et al. 2018), we broaden this approach by assessing both tolerance and lethality. By recapitulating a range of disease phenotypes and performing transcriptomic analysis, we directly identified differences in the host response that lead to divergent outcomes. To determine whether sex influences disease progression, we disaggregated our data by sex. This represents the first systematic analysis of sex as a biological variable in MERS-CoV mouse infection, revealing that sex modulates both tolerant and lethal responses.

Risk factors are biological, environmental, social, or behavioral variables associated with an increased risk of severe illness, according to the WHO (WHO 2024a). In humans, sex is a known risk factor for severe MERS, yet age has been the only risk factor investigated in hDPP4 mice to date (Fan et al. 2018; Algaissi et al. 2019; Iwata-Yoshikawa et al. 2019). Historically, sex as a biological variable has received little investigation; only after funding agencies mandated its consideration did the inclusion of both sexes become more routine (NIH 2015; Service Canada 2014). Despite these policy changes, a 2020 bibliometric analysis found that few studies disaggregate and analyze data by sex (Woitowich, Beery, and Woodruff 2020), leaving the biological impact of sex underexplored. In this study, we investigated how sex influences early responses to MERS-CoV infection in hDPP4 mice exposed to either a low or high dose.

We show that male mice succumb to infection at both doses (Fig. 5B), consistent with epidemiological data that males are more susceptible to severe MERS (Alghamdi et al. 2014). In contrast, disease outcomes in females vary by dose: high dose infection is uniformly lethal, while 2 of 3 low dose mice survived (Fig. 5B), although one mouse did not seroconvert (data not shown).

To interrogate the mechanisms underlying these observed sex differences to MERS-CoV infection, we analyzed transcriptional responses both independent of and accounting for sex. DE analysis revealed a higher number of total DE genes in the lethal group compared to the tolerant group (Fig. 3A), resulting in distinct patterns of functional enrichment between dose groups (Fig. 3D). Interestingly, multidimensional scaling of all DE genes showed overlap between tolerant and lethal groups, but clear separation between sexes (Fig. 6B). This suggests that the magnitude of the transcriptional response is primarily driven by dose, while the nature of the response is shaped by sex. For example, lethal males had higher numbers of downregulated DE genes compared to tolerant males, whereas females showed minimal downregulation (Fig. 6A).

Specifically, males downregulated key antiviral molecules, such as IFNβ (Fig. 4C), and repressed multiple innate pathways. Such delayed antiviral responses have previously been associated with severe outcomes in MERS-CoV animal models (Li et al. 2016, 2017; Agrawal et al. 2015; Falzarano et al. 2014), as well as in experimental models of other highly pathogenic emerging viruses (Price et al. 2020; Rasmussen et al. 2014).

In contrast to males, females mount an immediate antiviral response following MERS-CoV infection. One clear example of this sex difference is the activation of PKR signaling. PKR is encoded by *EIF2AK2*, an IFN-inducible gene (Kerr, Brown, and Hovanessian 1977; Meurs et al. 1990), and is activated upon binding double-stranded RNA, a pathogen-associated molecular pattern (García et al. 2006; Levin, Petryshyn, and London 1980). Once activated, PKR inhibits both host and viral translation, triggers apoptosis and amplifies the IFN response (Cesaro and Michiels 2021). Due to its broad antiviral activity, many viruses have evolved strategies to evade PKR-mediated signalling (Cesaro and Michiels 2021), including MERS-CoV which expresses accessory protein 4a (p4A) to antagonize this pathway (Rabouw et al. 2016). Despite p4A activity, we show that high-dose females are still able to upregulate *EIF2AK2* early in infection (Fig. 4B), resulting in activation of PKR-mediated signaling and apoptosis (Fig. 6D). In contrast, PKR signalling is absent in males and low-dose females. While the mechanism underlying this response remains unclear, *EIF2AK2* is downstream of several E2-regulated proteins (Fig. 4B) and is induced by type I IFN, which is generally more robust in females (Dunn, Perry, and Klein 2024).

Sex-specific differences in the host response to MERS-CoV reflect the disparity in viral read counts between males and females (Fig. S7), as well as innate immune biases determined by genomic and sex hormone differences (Klein and Flanagan 2016). We observed greater predicted activation of upstream female sex hormones, along with stronger direct and indirect connectivity between β-estradiol and DE anti-viral and pro-viral genes, compared to male sex hormones (Fig. 4). In the lethal group, females exhibited greater transcriptional activation and mounted an earlier antiviral response (Fig. 6), but had greater predicted activation of estrogen upstream regulators than tolerant females. This aligns with findings in other models showing that tolerance is associated with strict transcriptional control (Soares, Teixeira, and Moita 2017). Males exhibited sex-specific signaling involving IL-10 (Fig. 6D), an anti-inflammatory cytokine influenced by androgen signaling (Torcia et al. 2012). Together, these findings suggest that sex hormones may play a key role in regulating the host response to MERS-CoV infection, though further studies are needed to confirm these mechanisms.

Overall, this study establishes foundational evidence that sex influences MERS-CoV outcomes, likely through differences in the timing and magnitude of early immune responses. These findings underscore the importance of sex-disaggregated analyses in virology and pathogenesis research and provide new directions for identifying sex-specific risk factors in infectious diseases.

### Limitations

This study provides new insights into the role of sex during early MERS-CoV infection in hDPP4 transgenic mice infected at either a high or low dose. While the original aim was to investigate differences between tolerant and severe disease outcomes, rather than sex, the analysis revealed sex-based differences in disease progression. However, limited sample sizes for male and female mice, along with the absence of sex-disaggregated viral titre and load data, constrain the strength of these findings. To better understand the influence of sex on MERS-CoV pathogenesis, future studies should include larger cohorts of male and female mice, assess sex hormone levels during infection, and challenge gonadectomized animals to directly test the hormonal contribution to disease outcomes. Additionally, hDPP4 transgenic mice have been associated with neurological symptoms (Agrawal et al. 2015; Li et al. 2016), even at low doses, an outcome rarely seen in human MERS-CoV cases (Arabi et al. 2015). Therefore, future studies may benefit from using a model that better recapitulates human MERS-CoV disease. Finally, to capture a more complete picture of the host response, future work should incorporate RNA-seq, viral titration, and immunohistochemistry across multiple tissues.

## Data and code availability

All raw sequencing data is available through NCBI Bioproject Accession PRJNA1276520. Code is available at https://github.com/rasmussen-lab/MERS-CoV-tolerance-sex-hDPP4-mice.

## Author Contributions

Conceptualization: VJM, NvD, ALR; Data curation: MRH, RS, NvD, ALR; Formal analysis: MRH, RS, VJM, NvD, ALR; Funding acquisition: MRH, VJM, ALR; Investigation: MRH, TB, PS, VJM, NvD, ALR; Methodology: VJM, NvD, ALR; Project administration: VJM, ALR; Resources – VJM, ALR; Software: MRH, RS; Supervision: VJM, NvD, ALR; Validation: MRH, ALR; Visualization: MRH, RS, ALR; Writing – original draft: MRH, VJM, NvD, ALR; Writing – review and editing: MRH, TB, RS, PS, VJM, NvD, ALR.

## Acknowledgements

We would like to acknowledge Adam Price, Nina Deoras, Diane McFadden, and Atsushi Okumura for their contributions to sample acquisition and data management. We are also deeply grateful to Heinz Feldmann and Michael Kirby for their thoughtful comments and suggestions about the study and the manuscript. We acknowledge that generative AI (ChatGPT and Gemini) were used to generate code and identify copy editing or grammatical errors. Generative AI was not used to create text or figures in the manuscript.

## Methods

### Biosafety Statement

All animal experiments were approved by the Institutional Animal Care and Use Committee (IACUC) of Rocky Mountain Laboratories (RML). Experiments were performed according to the guidelines of the Association for Assessment and Accreditation of Laboratory Animal Care (AAALAC) and were carried out by certified staff in an AAALAC-approved facility. Experiments were conducted in a biosafety level 4 (BSL-4) laboratory at RML with approval from the RML Institutional Biosafety Committee (IBC). Animals were group housed. The animal room was climate-controlled with a fixed light-dark cycle of 12-hour light/12-hour dark. Food and water was available *ad libitum.* Animals were monitored at least once daily, and twice daily upon showing signs of disease. Animals were anesthetized via isoflurane and euthanized via cervical dislocation when they reached endpoint criteria as determined by the RML IACUC approved score parameters.

### Cells and virus culture and titration

MERS-CoV-EMC/2012, also known as *Betacoronavirus cameli*, was kindly provided by Erasmus Medical Center, Rotterdam, the Netherlands. Virus stocks were produced in VeroE6 cells grown in DMEM (Sigma) supplemented with 2% fetal calf serum (Logan), 1 mM L-glutamine (Lonza), 50 U/ml penicillin and 50 μg/ml streptomycin (Gibco). VeroE6 cells were maintained in DMEM supplemented with 10% fetal calf serum, 1 mM L glutamine, 50 U/ml penicillin and 50 μg/ml streptomycin. MERS-CoV was titrated in VeroE6 cells with tenfold serial dilutions of virus in 2% DMEM. Cells are incubated at 37°C and 5% CO2 for five days, after which cytopathic effect was scored and TCID_50_ was calculated from four replicates using the method of Spearman-Karber. For tissue titrations, tissue was homogenized, in 2% DMEM, spun down for 10 min at 8000 rcf, and supernatant was titrated.

### Animal experiments

To simulate differential outcomes to infection, male and female hDPP4 mice (van Doremalen et al. 2024) were infected with either a low (tolerant) or high (lethal) infectious MERS-CoV dose diluted in sterile DMEM. Tolerant and lethal groups, as well as time-matched mock-infected controls were anesthetized with inhalation isoflurane and infected via intranasal inoculation using a total volume of 25 µl. Infected and time-matched controls were serially euthanized at days 1, 2, 3, 4 and 5 post infection (N=6 per time point per dose, consisting of 2-4 females or males); lung samples were collected, frozen and later used for RNA extraction and titration. A secondary group (N=6 per dose) was observed for survival for up to 28 days.

### RNA extraction and QC

Tissues were weighed and homogenized in RLT buffer (Qiagen) and RNA was extracted using the RNeasy method (Qiagen) according to the manufacturer’s instructions. Viral RNA detection was performed via probe-based one-step real-time qPCR: 5 μl RNA was used in the Rotor-GeneTM probe kit (Qiagen) according to instructions of the manufacturer. Standard dilutions of a virus stock with known titer were run in parallel in each run, to calculate TCID_50_ equivalents in the samples. For viral RNA detection, we targeted upstream of the envelope gene sequence (UpE) (ref Corman, V. M. et al. Detection of 2019 novel coronavirus (2019-nCoV) by real-time RT-PCR. Eurosurveillance 25, 2000045 (2020)). MERS-CoV M-gene mRNA was detected as described by Coleman et al. (Coleman, C. M. & Frieman, M. B. Growth and Quantification of MERS-CoV Infection. Curr. Protoc. Microbiol. 37, 15E.2.1–9 (2015)).

### mRNA sequencing

Total RNA samples were poly-A enriched and underwent library preparation using Illumina TruSeq v2 reagents. Libraries were run on an Illumina HiSeq 4000 short read sequencer at the Icahn School of Medicine at Mt. Sinai Genomics Core Facility. For each sample, we collected a minimum of 30 million single-end 100 base pair (bp) reads.

### Data processing and normalization

Raw RNA-seq data was processed using FastQC v0.12.1, to remove adapters and assess read quality. To enable read mapping, a mouse genome index was built using the parameter “--sjdbOverhang 99”, from the *M. musculus* reference genome (GRCm39) and its corresponding annotation retrieved from the Ensembl database release 112. High quality reads were aligned to the assembled mouse genome using STAR v2.7.11b. Counts of the aligned reads were determined using the Rsubread v2.18.0 Bioconductor package in R v4.4.0. Low expression transcripts were excluded from differential expression analysis, according to the inclusion criteria of counts of 15 or higher in at least six samples. In addition, raw RNA-seq data was aligned to the MERS-CoV EMC 2012 genome (NCBI RefSeq accession GCF_000901155.1) to assess viral transcript counts.

### Differential expression analysis

Differential expression (DE) analysis was performed in the Bioconductor package DESeq2 v1.44.0 using the Wald test. DE analysis was performed twice. The first analysis identified genes differentially expressed in tolerant and lethal outcome groups relative to time-matched mock-infected controls. The second analysis identified DE genes relative to time-and sex-matched mock-infected controls. Transcripts from infected groups were considered to be differentially expressed if the Benjamini-Hochberg adjusted *p*-value was 0.05 or less and the fold change greater than |1.5| when compared to time-matched controls. The number of genes meeting DE criteria were visualized using R v4.4.0.

### Multidimensional scaling

To examine the similarity of gene expression profiles across samples, read counts underwent multidimensional scaling (MDS). MDS calculates the Euclidean distances between samples; Euclidean distance quantifies overall gene expression similarity between two samples, then projects this measurement into a 2D or 3D space for visualization. Samples that cluster have similar gene expression profiles, while those that separate are transcriptionally distinct. MDS scaling was conducted on variance-stabilized read counts to assess global gene expression across samples as well as DE genes specifically, to investigate differences in the host response. All MDS quantification was performed in R v4.4.0 using the ggplot2 package for visualization.

### Functional analysis

DE transcripts (fold change > |1.5|; adjusted p < 0.05) were uploaded to QIAGEN Ingenuity Pathway Analysis (IPA) and were subjected to IPA Core Analysis. To identify the enriched pathways associated with DE genes, the canonical pathways were investigated and compared between experimental groups and controls. In addition, upstream regulator analysis was performed to assess the predicted role of sex hormone signalling in the observed gene expression changes. For both canonical pathway and upstream regulator analyses, a *p*-value cut-off of <0.01 and an absolute Z-score value of 2 was applied. Furthermore, to assess the connection between temporal changes in the host response and sex hormone activity, DE molecules associated with antiviral signaling–interferon α/β and ISGylation pathways–and proviral calcium signalling pathways were used to generate an IPA custom network. The IPA Overlay tool was used to superimpose both experimentally derived and predicted gene expression values of DE molecules on days 2 and 4 post-infection. DE molecules with no connections to either sex hormone and less than 2 direct connections to other DE molecules were removed.

---

**Figure S1.**
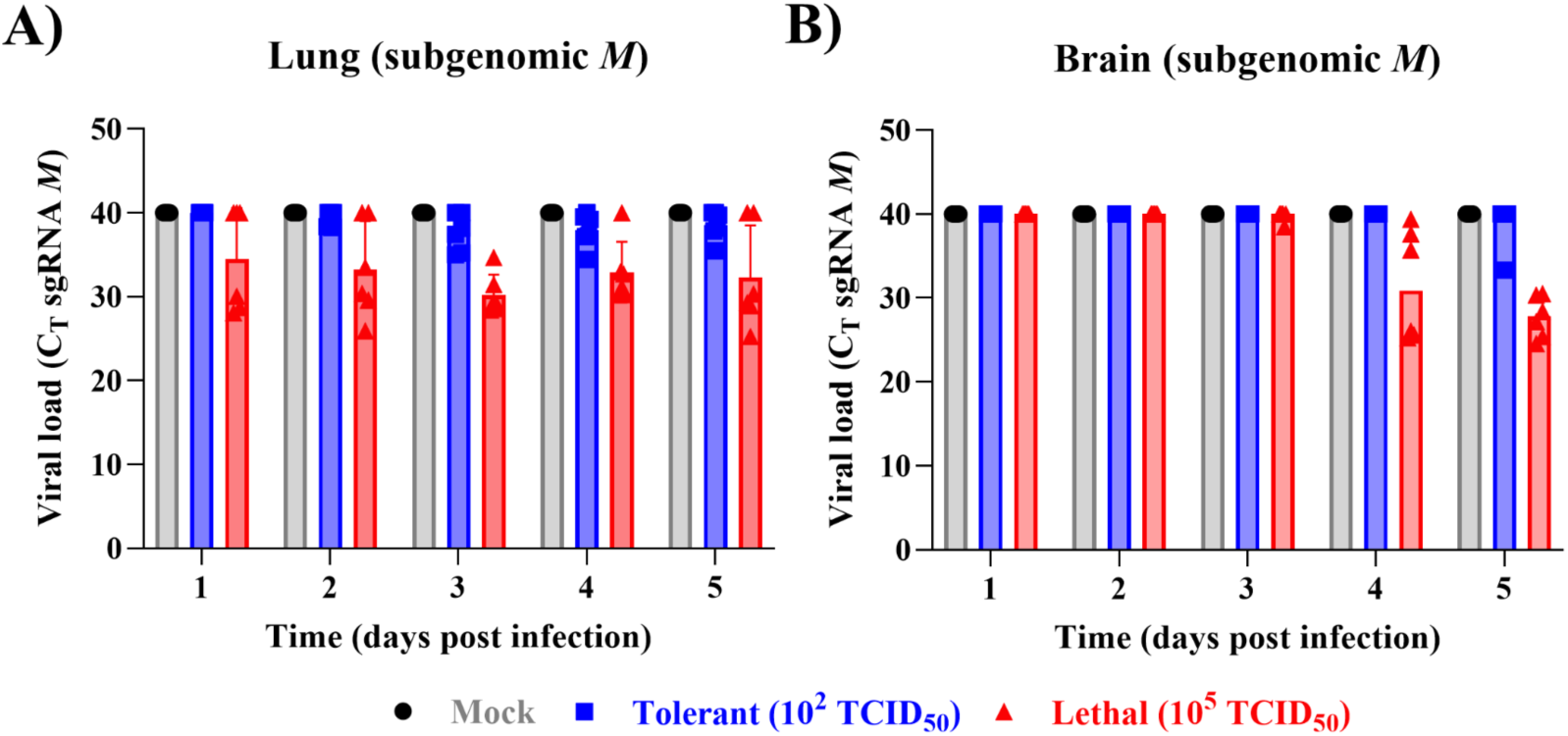
Mice infected with a high dose of MERS-CoV experience greater subgenomic replication in the lungs and brain compared to low dose mice. Lung (**A**) and brain (**B**) tissues were collected days 1-5 post-infection and RNA was extracted to assess viral load by RT-qPCR. The MERS-CoV subgenomic (sgRNA) *M* gene was used to assess subgenomic replication. Data represent the median for 6 mice per group.

**Figure S2.**
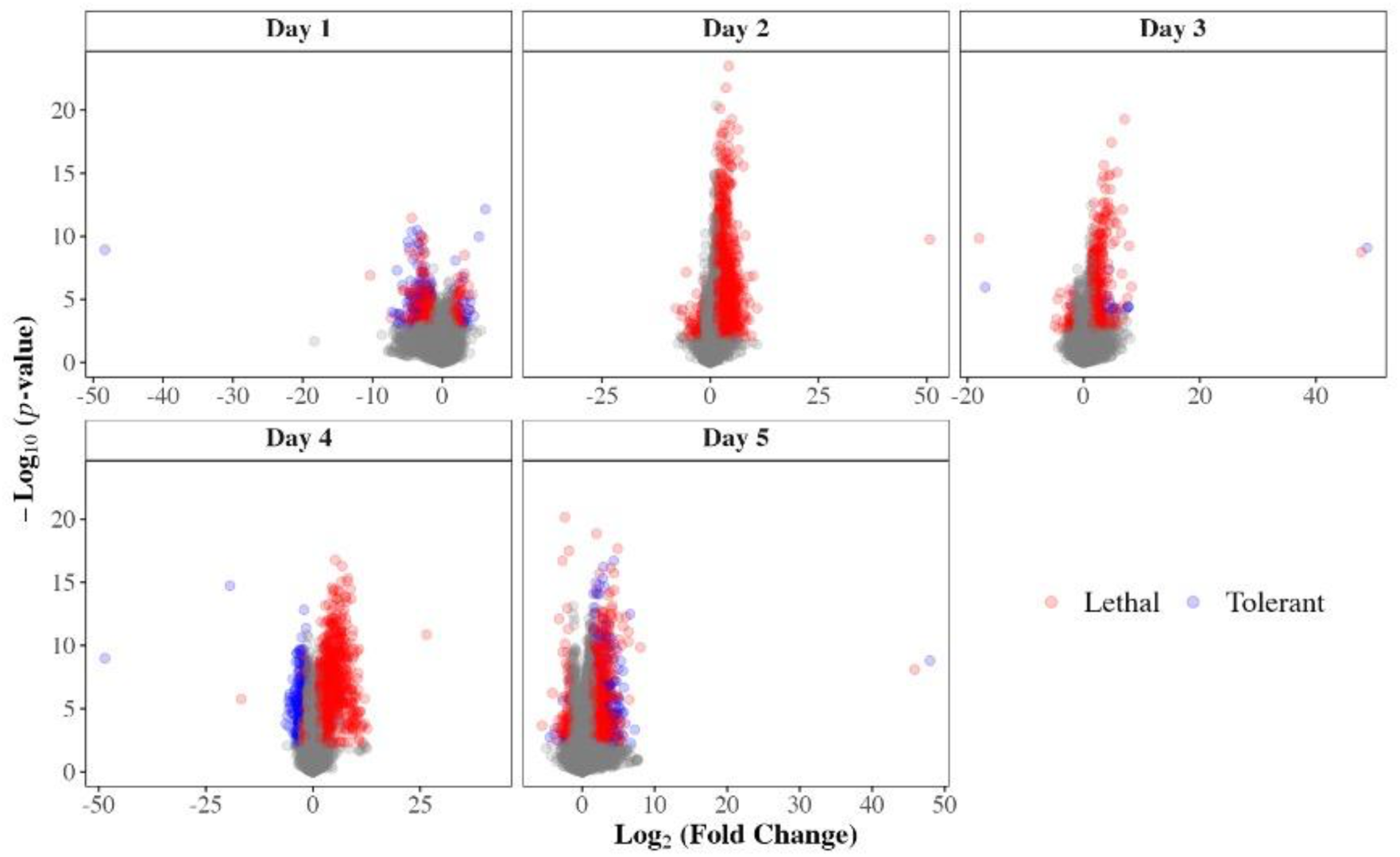
Increased differential gene expression in lethal hDPP4 mice compared to tolerant group. Lung samples were collected on days 1-5 post-infection from hDPP4 mice infected with a tolerant (10^2^ TCID_50_) or lethal (10^5^ TCID_50_) dose of MERS-CoV. Samples were subjected to RNA-seq followed by data processing and differential expression (DE) analysis relative to time-matched mock-infected controls. The number of DE genes (*p*<0.05, log2FC > |1.5|) in tolerant and lethal groups were compared. Lung samples were collected from 6 mice per timepoint per dose group.

**Figure S3.**
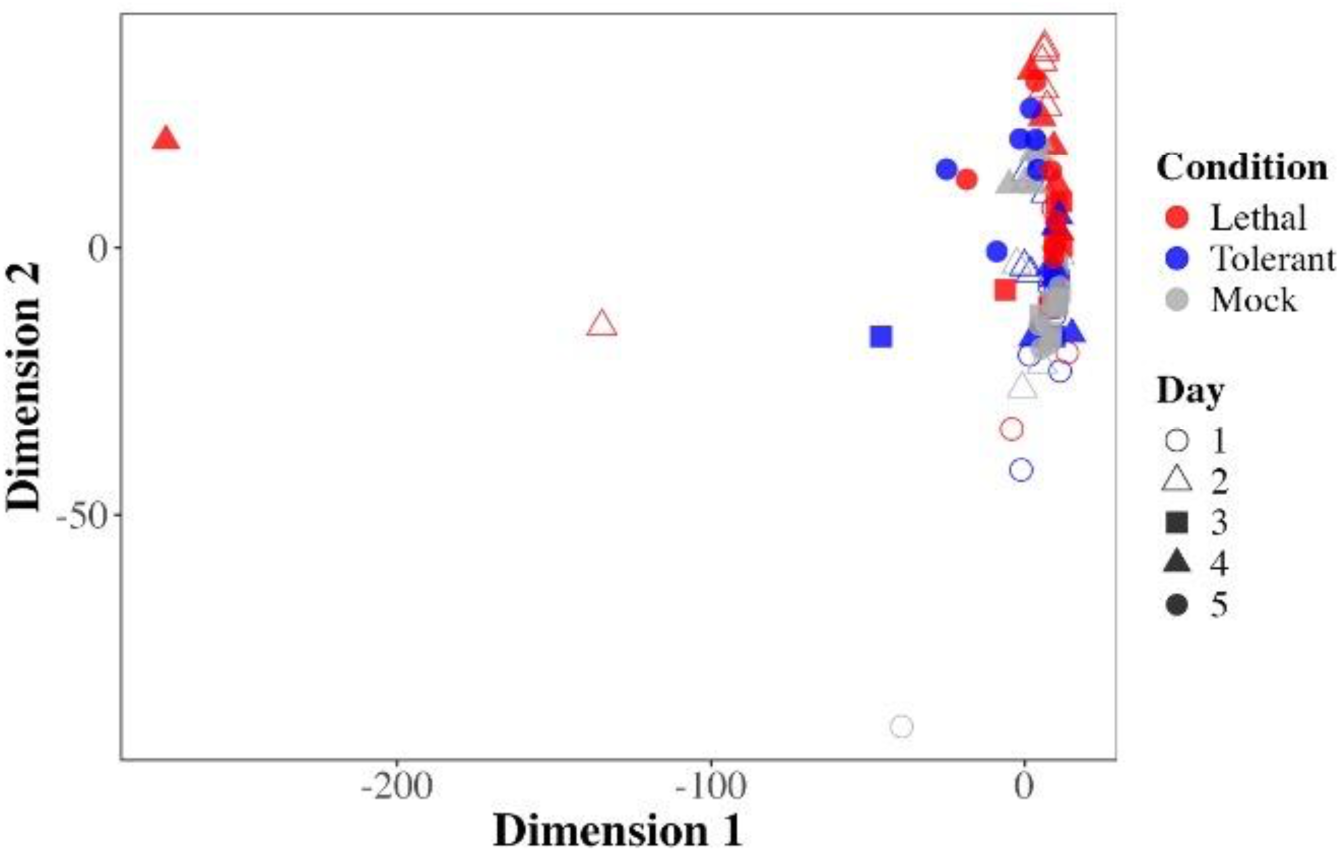
Normalized counts cluster together regardless of dose group. Lung samples from tolerant (10^2^ TCID_50_), lethal (10^5^ TCID_50_) and mock hDPP4 mice infected with MERS-CoV were collected days 1-5 post-infection and subjected to RNA-seq. Read counts were variance stabilized and subsequently subjected to Euclidean distance calculation to enable multidimensional scaling (MDS). Lung samples were collected from 6 mice per timepoint per dose group.

**Figure S4.**
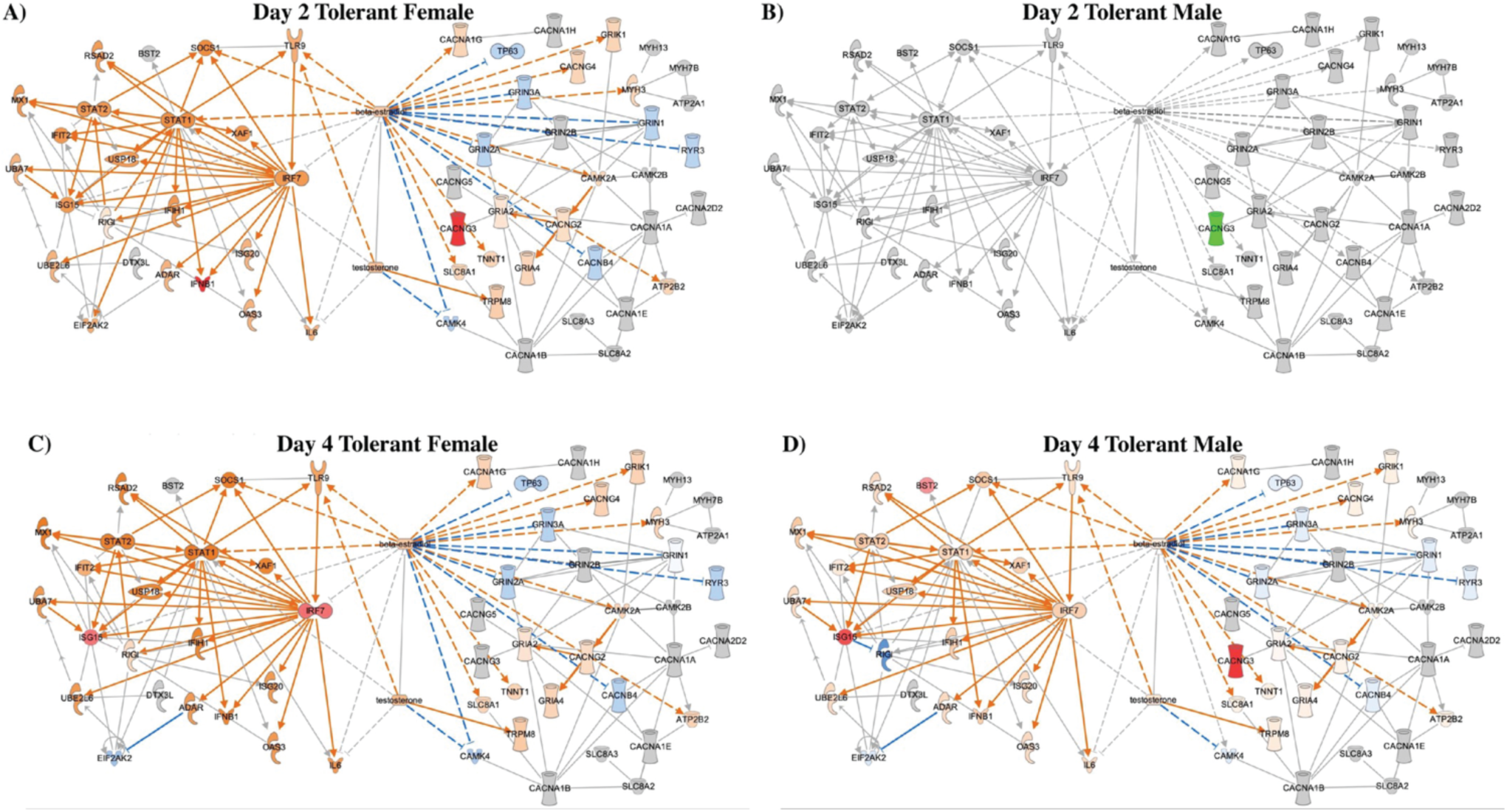
Tolerant female and male hDPP4 mice display temporal differences in their MERS-CoV infection response. Male and female mice were infected with a tolerance-simulating dose of 10² TCID_50_ or were mock-infected. RNA samples were collected from the lung on days 1-5 post-infection then underwent RNA-sequencing. Reads were subjected to differential expression (DE) analysis relative to time-matched mock-infected controls. DE genes (*p*<0.05, log2FC > |1.5|) were uploaded to Ingenuity Pathway Analysis (IPA) for custom network mapping (**A**-**D**). DE molecules from antiviral interferon α/β and ISGylation pathways (left) were connected and compared against connected DE molecules from the pro-viral calcium signaling pathway (right). The IPA Overlay tool was used to visualize expression in days 2 and 4 post-infection: red colouring indicated experimentally derived upregulation, while orange and blue represented predicted up or downregulation, respectively. Colour brightness indicated degree of enrichment.

**Figure S5.**
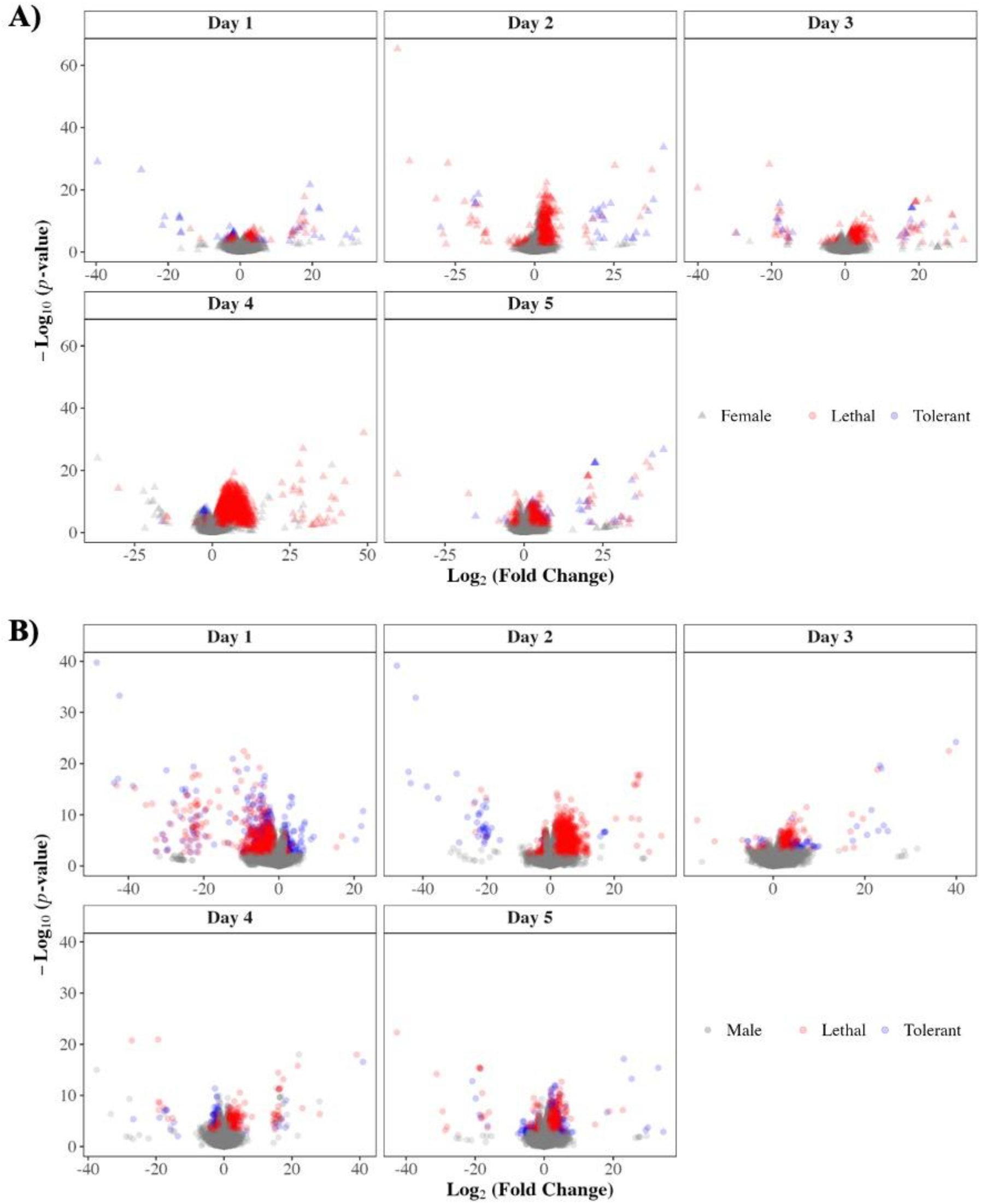
Female and male mice infected with MERS-CoV display dose and sex-dependent differences in DE of genes. hDPP4 mice were infected with a tolerance-inducing (10^2^ TCID_50_) or lethal (10^5^ TCID_50_) dose of MERS-CoV, or were mock-infected. On days 1-5 post-infection, lung samples were collected and subjected to RNA-seq. Reads underwent differential expression (DE) analysis by comparing lethal or tolerant gene expression to that of time-matched mock-infected controls. The number of DE genes (*p*<0.05, log2FC > |1.5|) in lethal and tolerant groups over time was compared between females (**A**) and males (**B**). Lung samples were collected from 6 mice per timepoint per dose group.

**Figure S6.**
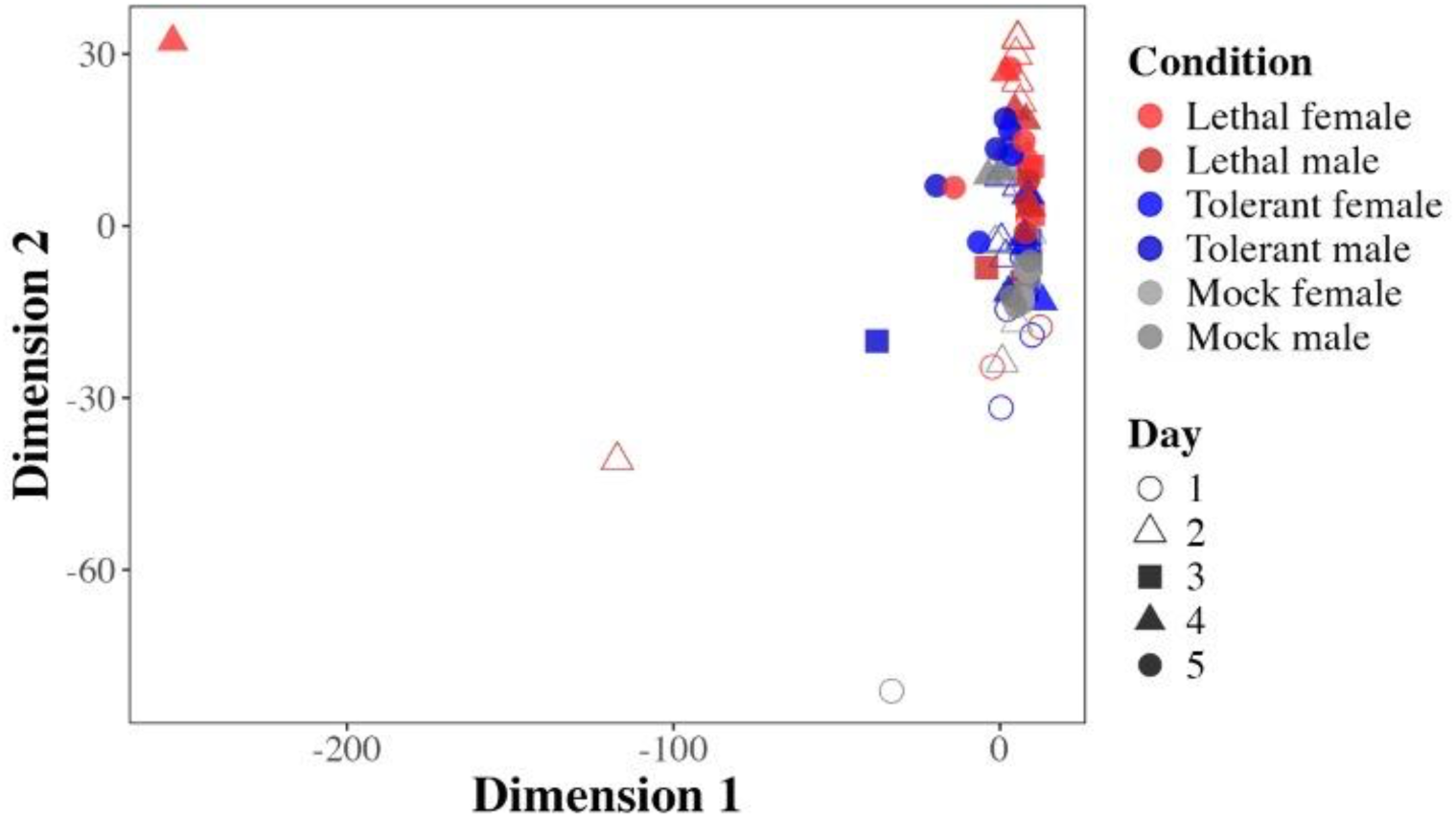
Normalized read counts cluster together regardless of dose group or sex. Lung samples from male and female tolerant (10^2^ TCID_50_), lethal (10^5^ TCID_50_) and mock hDPP4 mice infected with MERS-CoV were collected days 1-5 post-infection and subjected to RNA-seq. Read counts were variance stabilized and subsequently subjected to Euclidean distance calculation to enable multidimensional scaling (MDS). Lung samples were collected from 6 mice per timepoint per dose group.

**Figure S7.**
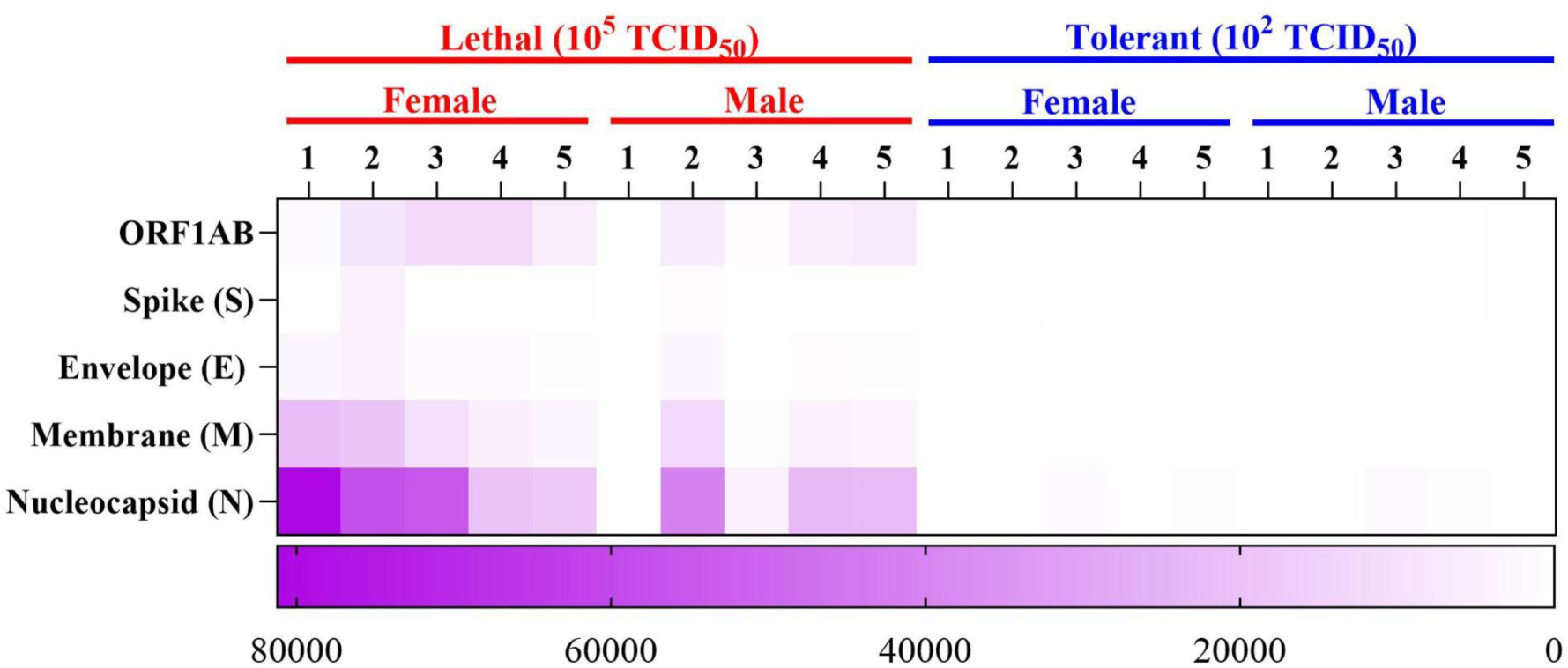
Increased MERS-CoV transcripts in lethal hDPP4 mice compared to tolerant group. Lung samples collected days 1-5 post-infection were collected and subjected to RNA-seq. Reads were processed and filtered to allow alignment of viral transcripts to the MERS-CoV EMC/2012 genome. Read counts across MERS-CoV transcripts were visualized.

